# ELEMENT-FREE MULTISCALE MODELING OF LARGE DEFORMATION BEHAVIOR OF RED BLOOD CELL MEMBRANE WITH MALARIA INFECTION

**DOI:** 10.1101/136648

**Authors:** A.S. Ademiloye, L.W. Zhang, K.M. Liew

## Abstract

In normal physiological and healthy conditions, red blood cells (RBCs) deform readily as they pass through the microcapillaries and the spleen. In this paper, we examine the effects of *Plasmodium falciparum* infection and maturation on the large deformation behavior of malaria-infected red blood cells (iRBCs) by means of a three-dimensional (3D) multiscale meshfree method. We numerically simulated the optical tweezers experiment and observed the force-displacement response of the iRBC membrane as malaria infection progresses. Our simulation results agree well with experimental data and confirm that the deformability of malaria-infected cells decreases significantly as malaria infection progresses.

## 1 INTRODUCTION

Malaria is the most important parasitic disease of humans and claims the lives of more children worldwide than any other infectious disease. The World Malaria Report 2015 published by the World Health Organization (WHO) revealed that, in the year 2015 alone, around 214 million new cases of malaria infection and ∼438,000 malaria deaths occurred worldwide [1]. Despite the huge progress recorded in the past few years, the recurrent development of antimalarial drug resistance remains a significant source of concern for all. Of all the known malaria parasites, *Plasmodium falciparum (P. falciparum)*, which is a protozoan parasite transmitted by the female Anopheles mosquito, has been identified as the most prevalent and lethal malaria parasite affecting humans [2]. After invading the healthy RBCs, the merozoites develop through the ring (*Pf*-rRBC), trophozoite (*Pf*-tRBC) and schizont (*Pf*-sRBC) stages with varying properties and degrees of influence on the cell properties. The product of this dynamic invasion process is the bursting of the RBCs resulting in anemia, chills, and fever [3]. Other notable effects of infection include increased membrane shear modulus, cell viscosity and cytoadherence [4].

Noteworthy numerical studies into the effect of malaria infection on RBC membrane include the works by Dupin et al. [5] using a 3D lattice Boltzmann model, Hosseini and Feng [6] by means of smoothed particle hydrodynamics (SPH) method and dissipative particle dynamics (DPD) method by Fedosov et al. [7]. In a bid to predict the large deformation behaviors of the iRBC membrane more precisely and elucidate their stiffening mechanism, a nonlinear 3D multiscale RBC membrane model is proposed in this study. The developed 3D multiscale RBC multiscale model, based on the 2D atomistic-continuum model presented in Ref. [8], is computationally efficient and able to capture the atomistic scale behavior of RBC membrane more accurately. This approach has been successfully applied to study the biomechanical behavior of healthy RBC membrane [9–12].

## 2 MULTISCALE MESHFREE COMPUTATIONAL FRAMEWORK

### 2.1 Multiscale hyperelastic constitutive model

In this section, we describe the development of a hyperelastic constitutive model that is derived from the first-order Cauchy–Born rule by using the coarse-grained Helmholtz free energy to describe the membrane atomic interactions. The Cauchy–Born rule establishes a connection between the deformation of the lattice vector of an atomistic system and that of a continuum displacement field, and plays an important role in the development of continuum constitutive models of atomic lattices. Considering a representative microstructure that is composed of six spectrin links *I-J*(*J*=1,…,6) as shown in Fig. 1 above, the deformation of each spectrin links can be approximated using the Cauchy-Born rule, as follows,

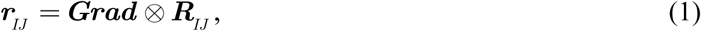

where ***Grad =*** *Grad*_*iJ*_*e*_*i*_⊗*e*_*J*_, ***R***_*IJ*_ and ***r***_*IJ*_ denotes the first-order deformation gradient tensor, the undeformed and deformed spectrin link length between junction complexes *I* and *J*, respectively.

**Figure 1:**
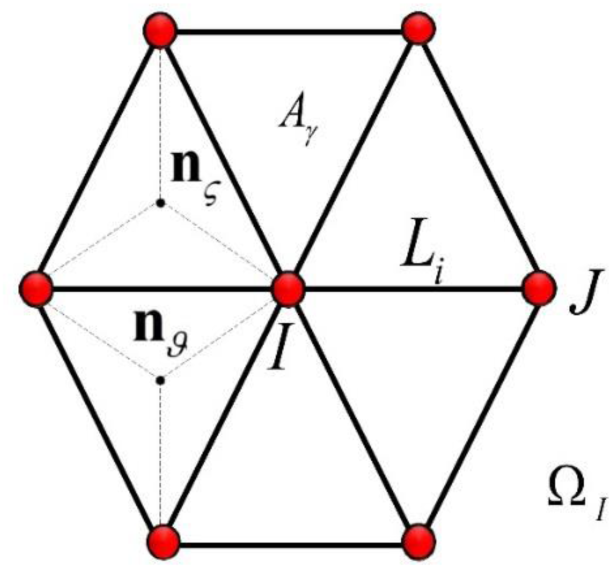
Representative microstructure of the RBC membrane

In our current derivation, the strain energy density function *W*_0_ at a junction complex *I* of the RBC membrane representative microstructure is defined using the coarse-grained Helmholtz free energy [8,9,13] at this point, which is obtained as a summation of membrane in-plane and bending energies, and expressed as

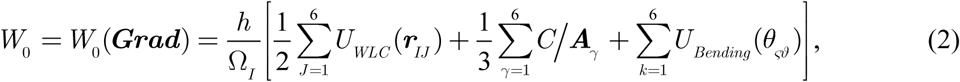

where *h* represents RBC membrane thickness and Ω*I* is the average area per junction complex in the reference configuration, calculated using 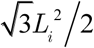. The first-order Piola-Kirchhoff stress tensor ***P*** and the tangent modulus matrix ***M*** corresponding to the first- and second-order derivative of the strain energy density with respect to ***Grad***, respectively can be calculated using,

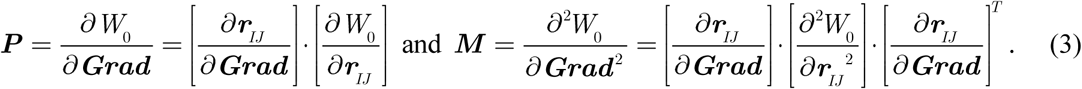

### 2.2 Nonlinear meshfree computational framework

A nonlinear three-dimensional (3D) multiscale meshfree method based on the improved moving least-squares (IMLS) approximation and Ritz minimization scheme, is employed to investigate the effect of malaria infection on the large deformation behavior of iRBC membrane. The current approach is well able to overcome the inherent limitations of mesh-based methods since it satisfies the higher order continuity requirements, circumvent mesh distortion, increased computation cost due to remeshing process and owing to the nonlocal intrinsic properties of meshfree methods, displacements are the only nodal degree of freedom. We refer readers who are interested in the derivation and numerical computation procedure of the expressions in (1)–(3) above as well as the meshfree development and implementation to the following literature and the references cited therein [8,9,14,15].

### 2.3 Numerical simulation procedure

The full biconcave geometry of the RBC membrane was used for all numerical simulations in this study since the precise shapes of the various infection stages are unknown. This geometry was first discretized with *n_m_* = 162 meshfree nodes (Figure 2) and 766 quadrilateral background cells with four (4) Gauss points in two directions. Stretching forces from 0-195 pN was applied to the ends of the cell and pulled. The variation of healthy and infected RBC membrane deformed axial *d*_*a*_ and transverse *d*_*t*_ diameters are computed and plotted against increasing stretching force. The sets of RBC membrane microstructure parameters presented in Table I are dependent on the condition of the cell membrane and we opined that the as malaria parasite develops, the equilibrium spectrin link length increases and the persistence length decreases. Other simulation parameters are defined irrespective of the RBC condition. They include the physiological temperature, *T=*300 K (27°C), membrane thickness, *h=*12 nm and the RBC membrane bending coefficient, 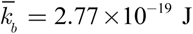 Results obtained from our numerical simulations are presented in comparison with experimental results reported in Ref. [16].

**Figure 2:**
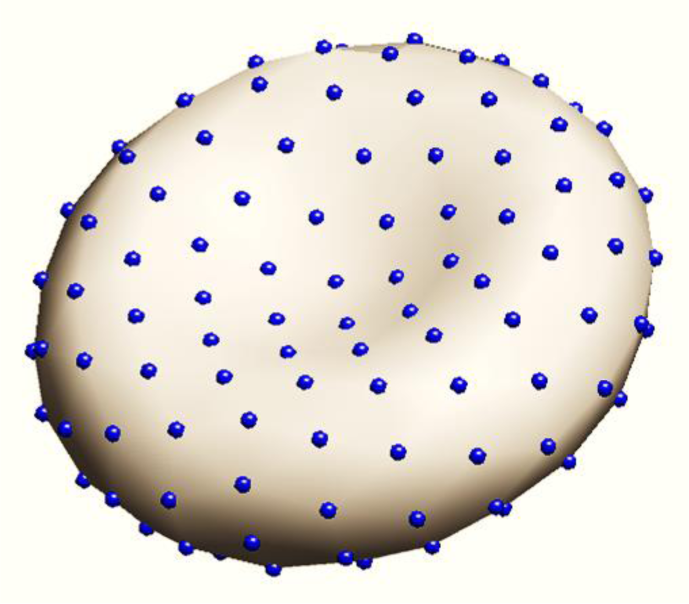
Discretization of iRBC membrane surface with 162 meshfree nodes

**Table I.**
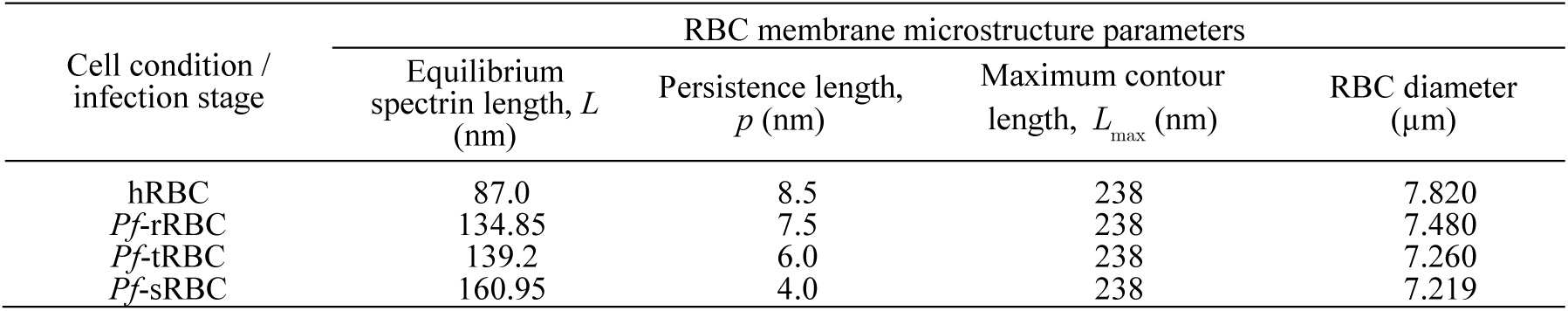
Values of microstructure parameters for healthy RBC and iRBC membrane

## 3 RESULTS AND CONCLUSIONS

Figure 3 shows the relationship between the variation in RBC membrane axial and transverse diameters and applied stretching force as malaria infection progresses in comparison with experimental data. The obtained result compares well with experimental result. We observe that as the malaria parasite develop, the deformability of the iRBC membrane decreases similar to the experiment reported in Ref. [16]. Our findings reveal that the increased stiffening of RBC membrane may be primarily due to the rearrangement and changes in the microstructure of the membrane underlying spectrin network rather than change in RBC membrane shape or formation of nanoscale knobs on the membrane surface.

**Figure 3:**
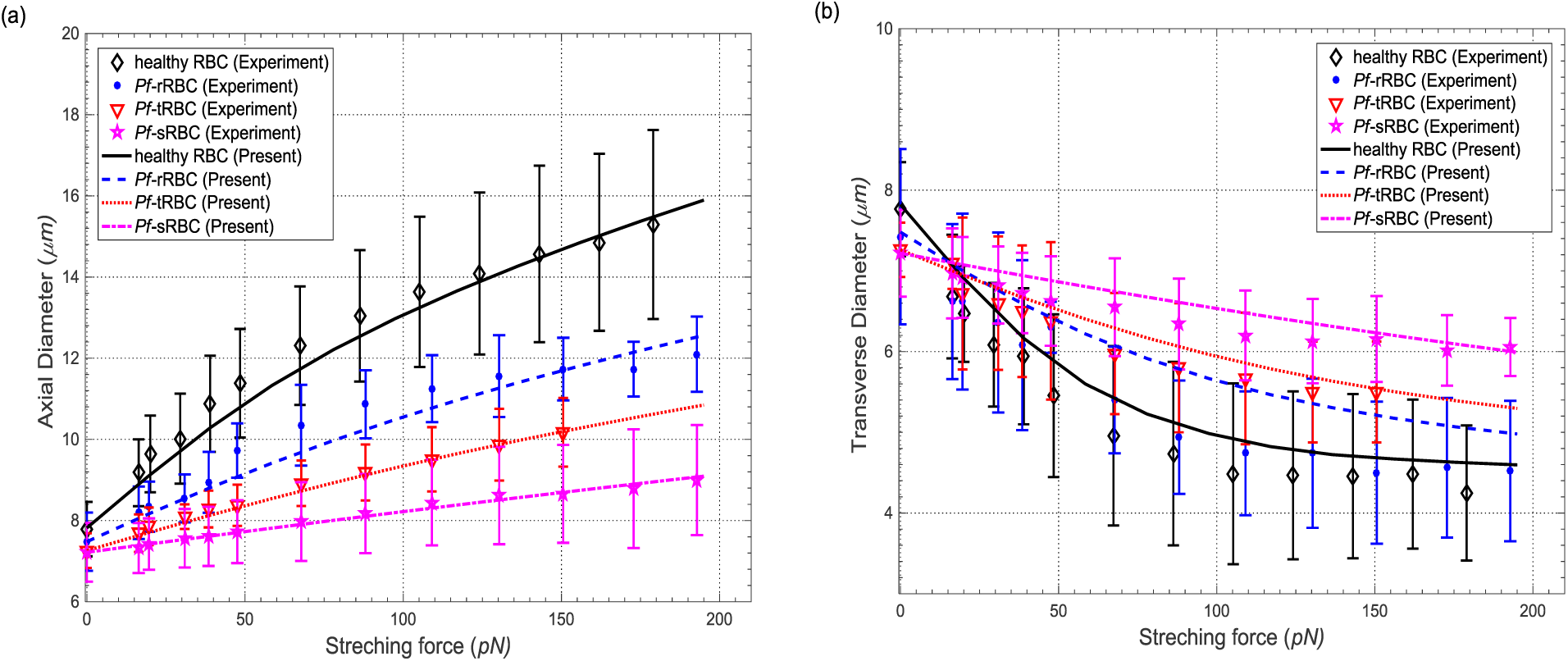
Variation of (a) axial diameter and (b) transverse diameters of healthy, *Pf*-rRBC, *Pf* tRBC and *Pf*-sRBC membrane in comparison with experimental data [16].

## ACKNOWLEDGEMENT

The authors gratefully appreciate the supports provided by Research Grants Council of the Hong Kong Special Administrative Region, China (Project No. 9042047, CityU 11208914) and National Natural Science Foundation of China (Grant No. 11402142 and Grant No. 51378448).

